# Evidence supporting resting-state power spectrum exponent as a non-invasive measure of neural-circuit excitation:inhibition balance in children with and without autism

**DOI:** 10.64898/2026.11.23.767072

**Authors:** J. Christopher Edgar, Mia Pearce, Yuhan Chen, Heather L. Green, Drayton L. Murray, Marybeth McNamee, Marissa DiPiero, Alice Onyango-Opiyo, Brianna Felgoise, Lisa Blaskey, Gregory A. Miller, Muhammad G. Saleh

## Abstract

**Introduction:** An ongoing neural circuit excitation (E) and inhibition (I) imbalance is thought to be a major contributor to clinical conditions such as autism, schizophrenia, and epilepsy. Current methods for directly measuring E:I dynamics require invasive procedures that are difficult to perform in non-human animals and nearly impossible in humans. A non-invasive measure of E:I balance is of great interest as a diagnostic as well as therapeutic marker, and as a means of studying relevant pathological and intervention mechanisms. Scientists have recently developed a computational model to show that, using resting-state (RS) MEG or EEG recordings, the E:I balance can be estimated from the power law exponent (slope) of the electrophysiological power spectrum. In their model, the RS power spectrum exponent is hypothesized to reflect the integration of synaptic currents in the brain, an index of E:I balance driven largely by excitatory (glutamate, Glu) and inhibitory (gamma-aminobutyric acid, GABA) neurotransmitter levels. Glu and GABA are the principal excitatory and inhibitory neurotransmitters in the brain, together shaping the balance between neural circuit excitation and inhibition. The present study shows that parietal-occipital neurochemistry as a predictor of the parietal-occipital RS exponent is driven by inhibitory neurotransmitter levels in typically developing (TD) children but not in children with autism spectrum disorder (ASD).

**Methods:** Midline parietal-occipital RS power spectra (MEG) and Glu and GABA+ concentration (MRS) were obtained from 129 children (TD N = 84, ASD N = 45) 10.2 to 17.9 years old, with follow-up measures in a subset of the children providing 148 MEG and MRS datasets. Analyses focused on RS neural function and Glu and GABA+ concentration in parietal-occipital cortex.

**Results:** *t*-tests showed no group differences between the parietal-occipital exponent, Glu, GABA+, or Glu/GABA+ ratio measures (*p*s > 0.21). In the full sample, older age was marginally associated with a lower exponent (*p* = 0.07), and older age associated with lower Glu concentration (*p* < 0.05). Age and GABA concentration or Glu/GABA+ associations were not observed (*p* > 0.05). There was no main effect of Glu or Glu x Group interaction with the exponent (*p*s > 0.14). Simple effect analyses of group interactions for GABA+ (*p* < 0.05) and ratio (*p*<0.05) showed a GABA+ and exponent association for TD (*p* = 0.01) but not ASD (*p* = 0.40) and a ratio and exponent association for TD (*p* = 0.01) but not ASD (*p* = 0.83).

**Discussion:** The brain chemistry and RS exponent associations suggest that a simple method for estimating E:I ratio reduces a major hurdle in understanding a currently difficult-to-measure, yet fundamental aspect of neural computation that has been proposed as central to clinical conditions such as ASD, epilepsy, and schizophrenia. Present findings indicate that the contribution of local brain chemistry to neural-circuit E:I balance differs for older children with and without ASD, with GABA inhibition presenting the more promising contributor to the E:I balance in TD children. Findings have the potential for high impact, including identifying neural markers for use in tracking treatment response.

## Introduction

A neural circuit excitation (E) and inhibition (I) imbalance is thought to be a major contributor to clinical conditions, such as autism, schizophrenia, and epilepsy (Gandal et al., 2010, 2012; McCormick & Contreras, 2001; Rao & Lowenstein, 2015; Rubenstein & Merzenich, 2003). Current methods for directly measuring E:I dynamics require invasive procedures that are difficult to perform in behaving animals, and nearly impossible in humans. A non-invasive measure of E:I balance is of great interest as a diagnostic marker as well as a therapeutic marker to assess the effect of treatment on brain activity.

Several groups have proposed that E:I balance can be obtained from resting-state (RS) neural activity measures (Chini et al., 2022; Gao et al., 2017; Lombardi et al., 2017). For example, Gao et al. (Gao et al., 2017) developed a computational model to show that, using RS electroencephalography (EEG) or magnetoencephalography (MEG) recordings, E:I balance can be estimated from the power law exponent (slope) characterizing the electrophysiological power spectrum. The RS power spectrum exponent is hypothesized to reflect the integration of synaptic currents in the brain, an index of E:I balance, this balance largely driven by excitatory glutamate (Glu) and inhibitory gamma-aminobutyric acid (GABA) neurotransmitter levels. Glu and GABA are principal excitatory and inhibitory neurotransmitters in the brain, together shaping the balance between neural circuit excitation and inhibition (e.g., (Buzsáki et al., 2007; Hu et al., 2014; Porges et al., 2021)).

EEG/MEG-based E:I model predictions have received some validation in non-human and human data. For example, Gao et al. (Gao et al., 2017) found that administration of general anesthetic in macaques resulted in changes to the exponent that tracked consciousness over time. Human studies observing a steeper (more negative) exponent during sleep and general anesthesia are also consistent with increased inhibition during these states (Colombo et al., 2019; Gao et al., 2017; Kozhemiako et al., 2022; Lendner et al., 2020; Schneider et al., 2022).

The recent hypothesis that E:I balance can be estimated from the RS power spectrum has had a substantial impact on research, with a rapid increase in the number of EEG and MEG studies reporting on the RS exponent in clinical populations. A recent paper (Donoghue, 2025) provided a systematic literature review that found 177 studies reporting on the RS exponent in clinical populations: EEG (118; 67%), MEG (24; 14%), deep brain stimulation (22; 12%), intracranial EEG (8; 5%) and implanted random noise stimulation (2; 1%). Studies were found across 38 clinical conditions (e.g., epilepsy, autism spectrum disorder (ASD), schizophrenia, stroke, Parkinson’s disease). Donoghue’s Table 2 shows that of the 177 papers, only 16 were published before 2020 (most of which assessed RS neural activity in epilepsy).

Donoghue et al. (Donoghue, 2025) found that a majority of the 177 studies (98/177; 55%) discussed the RS exponent as a possible biomarker, with E:I ratio the most common stated interpretation of the RS exponent (103/177; 58%). However, in most of the clinical conditions, the results were difficult to interpret. Donoghue noted that (bold added), “The consistency of results across disorders varies, with the most studied disorders of epilepsy and Parkinson’s having arguably the most consistency in their results, as well as a high degree of consistency in neurological disorders such as stroke and disorders of consciousness. **By comparison, psychiatric disorders appear to generally have less consistent results, consistent with the longstanding difficulty in identifying consistent biomarkers in psychiatric conditions.”**

Donoghue (Donoghue, 2025) considers several factors that might account for the variability of findings in psychiatric conditions such as ASD, schizophrenia, and ADHD: differences within and across studies in age/development stage, patient symptoms, marked heterogeneity within most psychiatric conditions, possible treatment effects, and between-study differences in the frequency range used to compute the aperiodic measures. Donoghue also considers the possibility of anatomical specificity in group differences, this prospect not addressed in studies that compute the exponent measure via averaging across EEG or MEG sensors (An et al., 2025; Cazares et al., 2024; Chung et al., 2024; Dede et al., 2025; Webb et al., 2023).

Obtaining RS measures via averaging across EEG/MEG sensors is problematic for many reasons. As detailed in Edgar et al. (Edgar et al., 2026), noninvasive EEG and MEG scalp channels are too often interpreted as if a given sensor largely reflects neural activity near that sensor, such as a left anterior EEG electrode measuring left frontal lobe neural activity. Schaworonkow & Nikulin (Schaworonkow & Nikulin, 2022) showed that this is generally not the case. Assessing the contribution to each EEG sensor of RS alpha neural activity (8 to 12 Hz in adults) from different brain regions, they showed that each sensor reflects activity from across the brain. Given regional differences in the aperiodic offset and exponent measures as well as in their developmental maturation (Green et al., 2025; McNamee et al., 2026; Vandewouw et al., 2024), the best practice is to assess regional activity rather than average across sensors or brain regions (Edgar et al., 2026).

To make progress understanding the RS exponent, studies are needed that assess RS brain activity in region-specific source space rather than sensor space. With respect to validating the RS exponent as a measure E:I balance, also needed are studies assessing associations between the RS exponent and brain chemistry, most importantly Glu and GABA. To that end, the present study evaluated brain chemistry (MRS) and RS measures (MEG). Glu and GABA concentrations and RS exponent measures were obtained for a midline occipital/parietal region. In addition to being a brain area that provides MRS measures with excellent SNR, this region is of interest because dominant RS oscillation generators are concentrated in midline parietal-occipital areas (Berger, 1929; Edgar et al., 2022; Haegens et al., 2014; Huang et al., 2014; Salmelin & Hari, 1994).

Parietal-occipital RS aperiodic measures and parietal-occipital MRS Glu and GABA concentration measures were obtained in children with and without ASD^1^ 10 to 17 years old. Findings of an association between the RS exponent and MRS Glu and/or GABA+ would provide support for the hypothesis that the RS exponent provides a non-invasive assessment of neural-circuit E:I balance. Assessment of exponent and brain chemistry associations in children with ASD is of particular interest, as a prominent theory of brain dysfunction in ASD hypothesizes that the clinical presentation of ASD is substantially due to altered excitatory and inhibitory neurotransmission. This hypothesis, proposed by Rubenstein & Merzenich (Rubenstein & Merzenich, 2003), is based on the observation that many of the behaviors observed in ASD are consistent with a neural E:I imbalance. Postmortem studies have reported fewer sites where benzodiazepine could bind to GABA receptors as well as fewer GABA receptors in autistic brains (Blatt et al., 2001) and fewer GABA-mediated fibers in minicolumns in ASD (Casanova et al., 2003; Fatemi et al., 2014; Fetit et al., 2021; Yao & Li, 2024). Such findings led to hypotheses of too little GABA inhibition, also supported by studies showing a link between epilepsy (characterized by too little inhibition) and ASD (Hara, 2007; Jeste & Tuchman, 2015; Tuchman & Rapin, 2002). Later post-mortem ASD studies found differences in cortical inhibitory architecture, including alterations in GABA_A_ receptor expression, and interneuron density (Fatemi et al., 2009, 2014; Oblak et al., 2009; Yao & Li, 2024).

Genetic studies have consistently found that the genes associated with ASD operate on protein-protein interaction networks involved in synaptic functioning and chromatin remodeling (Autism Spectrum Disorder Working Group of the Psychiatric Genomics Consortium et al., 2019; Geschwind, 2011; Gilman et al., 2011). In the largest ASD exome sequencing study to date (35,584 individuals, including 11,986 with ASD), 102 ASD risk genes were identified (Satterstrom et al., 2020). Most converged on pathways involving the regulation of gene expression and neural communication. For example, GRIN2B, SHANK3, SYNGAP1, and SCN2A are essential for synaptic transmission, plasticity, and network excitability. The authors concluded (emphasis added): “In cells from the human cortex, expression of risk genes is enriched in excitatory and inhibitory neuronal lineages, consistent with multiple paths to an excitatory-inhibitory imbalance underlying ASD**…ASD genes provide compelling support for an excitatory-inhibitory imbalance in ASD through direct effects on neurotransmission.**”

Findings from a recent study using human cerebral cortex organoid models to assess the developmental consequences of ASD risk genes KMT5B, ARID1B, and CHD8 found that each gene was associated with asynchronous development of GABA-ergic interneurons and deep-layer excitatory projection neurons, with all three genes associated with abnormal neural-circuit activity during early stages of brain development (Paulsen et al., 2022). Finally, a recent review (Treccarichi et al., 2026) noted that genetic studies have identified ASD-associated variants affecting glutamatergic and GABAergic systems, as well as genes operating at their interface. As an example, synaptic adhesion and scaffolding genes such as NRXN1 and CNTNAP2 influence the formation and maintenance of excitatory and inhibitory synapses, with pathogenic variants in NRXN1 associated with ASD (Cooper et al., 2024; Onay et al., 2016). Treccarichi et al. (Treccarichi et al., 2026) concluded (emphasis added): “…findings support a model in which the glutamatergic and GABAergic pathways, while molecularly distinct, converge on shared developmental and functional processes regulating neuronal network stability…. **This convergence places E:I imbalance at the core of ASD pathophysiology and provides a unifying framework linking genetic susceptibility, synaptic dysfunction, and neurodevelopmental outcomes** (p. 21)”.

Present hypotheses were: (1) given the proposed Glu/GABA imbalance in ASD, TD and ASD children would differ in Glu and GABA concentration as well as the RS exponent, (2) findings of an association between the RS exponent and GABA (and perhaps Glu) in TD children would provide support for the RS exponent as a non-invasive measure of neural-circuit E:I balance, and (3) a RS exponent / GABA association would be observed in children with ASD.

## 2. Method

This study was approved by the CHOP Institutional Review Board, and all families gave written informed consent. When competent to do so, children 7 years of age and older gave verbal and written assent to participate. Families were compensated for their participation in the study.

### 2.1 Participants

MEG RS data were obtained from children participating in a longitudinal study examining the maturation of brain function and structure. Children were enrolled in the study between 10 and 15 years old, with longitudinal MEG RS and MRS brain chemistry measures obtained ∼1.5 and ∼3.0 years after the initial exam.

Edgar et al. (Edgar et al., 2023) provided details of how an ASD diagnosis was confirmed in ASD children and how study exclusion and inclusion criteria were confirmed in all children. All TD and ASD children met the following exclusion/inclusion criteria: (a) English as primary language, (b) no history of traumatic brain injury or other significant medical or neurological abnormality, (c) no premature birth (< 34 weeks gestation), (d) no uncorrectable sensory impairments (somatosensory, hearing, visual), (e) no sMRI contraindications, (f) no genetic conditions with a very high incidence of ASD (e.g., 22q/Fragile X), and (g) no neuroleptic/antipsychotic medication. At the time of the MEG+MRS exams, 10 of the children with ASD were prescribed stimulant medications. Nine of the children with ASD were taking antidepressants, one child with ASD was taking an anxiolytic, and six children with ASD were taking a non-stimulant medication for ADHD.

### 2.2 ASD Symptom and IQ Measures

Estimates of cognitive ability were obtained for: Full Scale IQ (measured via a 4-subtest short-form of the Wechsler Intelligence Scale for Children - Fifth Edition (WISC-V; (Wechsler, 2014)) or Wechsler Abbreviated Scale of Intelligence - Second Edition (WASI-II; (Wechsler, 2011)), estimated nonverbal IQ (eNVIQ; obtained from the Fluid Reasoning Index of the WISC-V or the Perceptual Reasoning Index of the WASI-II), and estimated verbal IQ (eVIQ; obtained from the Verbal Comprehension Index of the WISC-V or WASI-II). A measure of autism dimensional symptom severity, per parent report, was obtained using the Social Responsiveness Scale-2 (SRS-2) (Constantino & Gruber, 2012).

### 2.3 MEG Data Acquisition

Edgar et al. (Edgar et al., 2023) provided details of MEG and MRI data collection. In brief, MEG data were obtained using a 275-channel system (VSM MedTech Inc., Coquitlam, BC). To facilitate co-registration of MEG data to each child’s T1 MRI (see below), for each child three anatomical landmarks (nasion, right and left preauriculars), head position indicator locations, and scalp and face surface points were digitized using a Polhemus FastScan (Polhemus, Colchester, VT).

For the RS exam, over a 7 min period, children were periodically instructed to close their eyes, receiving instructions through insert earphones to open their eyes for 10 s approximately every two minutes. If the child opened their eyes during the exam, monitored by EOG, they were reminded to close them. Excluding three 10 s eyes-open segments, a total of 390 s of RS data were collected.

### 2.4 MEG Data Processing

MEG and sMRI data were co-registered using BESA 7.1 and BESA MRI 3.0 (MEGIS Software GmbH, Gräfelfing, Germany). MEG data were processed using BESA Research 7.1. A multi-step process was employed for removal of muscle and movement artifact. First, the child’s raw EOG and MEG data were visually examined. For the eyes closed (EC) exam, artifact correction was applied to remove heart-beat activity (Berg & Scherg, 1994; Scherg & Ebersole, 1993). Epochs with blinks, saccades, or other significant EOG activity were manually marked as artifacts. Review of the data for artifact was done blind to group status. A minimum of 60 s of artifact-free RS data was required for a child to be included in the analyses. Of the 148 RS datasets, 98% had > 150 s of RS data (see additional details in Section 3.1).

The following pipeline was applied to the artifact-free EC data to obtain measures of brain activity in source space. A standard source model was applied (the BESA BRM_Brains Regions_MEG source montage), projecting each child’s 275-channel MEG sensor data into brain source space, where the waveforms are the modeled source activities. The source montage uses 15 regional source locations covering lateral and midline frontal cortex, central and parietal cortex, and midline fronto-polar and occipito-polar cortex as well as bilateral anterior and posterior temporal lobes. Given two orthogonal dipoles per regional source, there are two time series at each location (Hoechstetter et al., 2004). These regional sources are not intended to correspond to individual neuroanatomical structures but rather to represent neural activity at coarsely defined regions and to provide measures of brain activity with better signal separation and with a higher signal-to-noise ratio than the sensor level provides (Berg & Scherg, 1994; Scherg & Ebersole, 1993). Locations of the regional sources in the model are such that there is an approximately equal distance between adjacent sources (3 cm), helping to separate signals originating from different brain regions. Given a parietal-occipital placement of the MRS voxel (see following text), the present study reports RS measures from the MEG-derived midline parietal source.

To transform MEG data from the time domain to the frequency domain, a Fast Fourier Transform was applied to artifact-free 3.41 s epochs of continuous data for each of the two orthogonally oriented time series at each regional source. Each epoch overlapped 50% with the next epoch, and each epoch was multiplied by a cosine window. This combination of overlap and windowing ensured that each time point contributed equally to the mean power spectra. When the segment of data following an artifact-free epoch was bad, there was no overlap between epochs, and windowing was thus applied to the next available artifact-free 3.41 s epoch. The mean power spectra for the two orthogonally oriented time series at each regional source were summed to yield power at a given frequency at the source.

### 2.5 Specparam Analyses of Source-Localized RS spectra

The specparam algorithm was used to parameterize the source-space RS spectra (Donoghue, Haller, et al., 2020; Ostlund et al., 2022a). Each power spectrum is treated as a linear combination of aperiodic activity and oscillatory peaks (individually modeled with Gaussians) that rise above the aperiodic signal. As detailed in specparam method papers (Donoghue, Dominguez, et al., 2020; Donoghue, Haller, et al., 2020; Ostlund et al., 2022a), outputs of the specparam algorithm include the offset and exponent values of the aperiodic signal and the bandwidth, power, and center frequencies of the identified oscillatory peaks. The following specparam fitting parameters were used: peak width limit 1 to 8 Hz; max number of peaks 3; minimum peak height 0.1; proximity threshold 2; and aperiodic mode fixed. Regression R^2^ and mean absolute error (MAE) measures, respectively representing the explained variance and total error of the model fit, served as goodness-of-fit measures used to identify potential outliers.

Available studies have not evaluated the appropriate frequency ranges for assessing the relationships proposed between specparam parameters and the MRS brain chemistry measures. The Gao et al. (Gao et al., 2017) brain chemistry and E:I association simulations assessed exponent values across a 20 to 150 Hz range (20 Hz window around the center frequency). Their Figure 1G plot shows stronger associations between the Glu/GABA E:I ratio and the power spectrum exponent at 20 Hz than for higher frequencies (exponent values below 20 Hz not assessed). Given almost no work in this area, present analyses examined associations between the parietal-occipital MRS GABA+ concentration and parietal-occipital exponent in several frequency ranges. In particular, the exponent was computed for a 1 to 55 Hz range (55 Hz the upper limit given power line noise at 60 Hz) and then the lower frequency increasing in 1 Hz steps up to 10 Hz (i.e., stopping at 10 to 55 Hz). For comparison with Gao et al., the exponent was also estimated using a 10 to 20 Hz, 20 to 40 Hz, and 30 to 50 Hz range.

**Figure 1.**
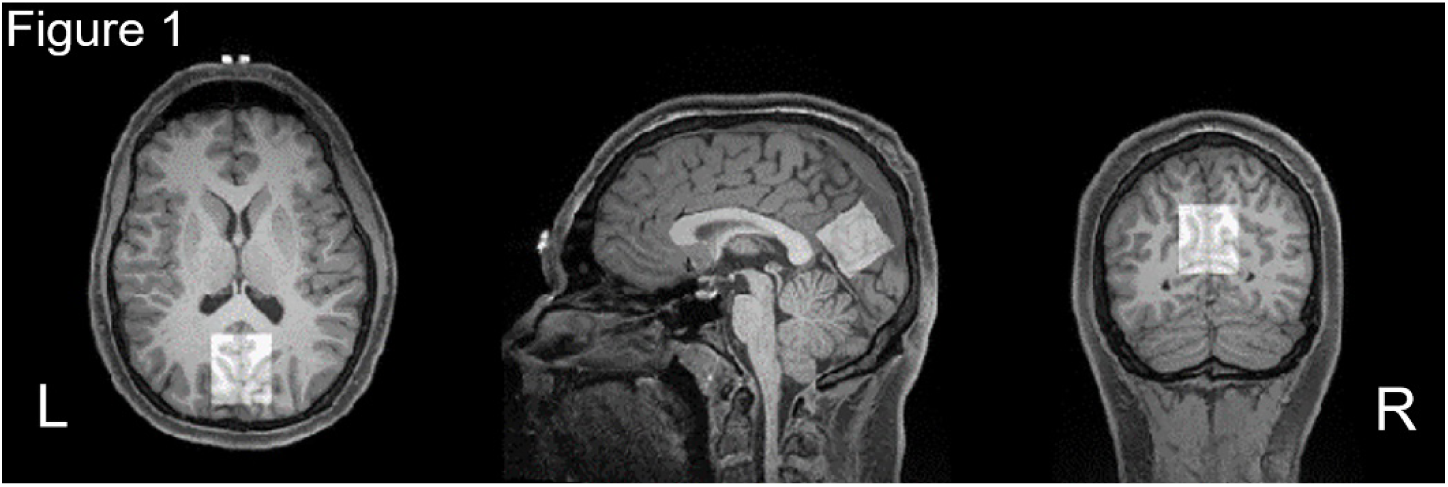
MRS voxel centered at midline of parietal-occipital cortex.

### 2.6 MRI and MRS Data Acquisition

sMRI provided T1-weighted, 3-D MP-RAGE anatomical images with 0.8x0.8x0.8 mm^3^ isotropic resolution (TR/TE = 2400/2.24 ms, flip angle = 8 degrees, and scan time ∼7 min ) for MEG source localization and MRS voxel localization at the midline of the parietal-occipital cortex and rotated in the sagittal plane to align the center of the voxel with the parietal-occipital sulcus (Figure 1). The HERMES protocol was then used to acquire GABA (Saleh et al., 2016, 2019) using the following parameters: 3x3x3 cm^3^ voxel size, TR/TE=2000/80 ms, 20 ms editing pulses, 224 averages, 2048 datapoints, and 2 kHz bandwidth. Water (unsuppressed) data were also acquired using the same parameters, except for 8 transients. The unedited PRESS acquisition was used to acquire Glu using the same parameters as the HERMES protocol, although with 32 averages (unsuppressed 8 averages) and a TE of 35 ms. The total scan time for HERMES and PRESS was ∼10 min.

### 2.7 Analysis of MRS Data

MRS data were processed on Osprey (Oeltzschner et al., 2020) to generate HERMES-based GABA-edited spectrum and the unedited Glu-edited spectrum from PRESS. Figure 2 shows typical spectra acquired from our cohort along with linear combination modeling for both spectra using LCModel (Provencher, 2001). To obtain water-scaled GABA (from the edited spectrum) and Glu (from the unedited spectrum) concentrations, we followed the procedures outlined in Saleh et al. (Saleh et al., 2016, 2019). Each participant’s T1 image was segmented in Osprey to calculate gray matter (GM), white matter (WM), and cerebrospinal fluid (CSF) voxel tissue fractions. Glu and GABA concentrations were corrected for partial volume and relaxation effects in each voxel using the method described by Gasparovic and colleagues ((Gasparovic et al., 2018); see equation in *OspreyQuantify*). Since the GABA signal at 3 ppm in the difference spectrum contains contributions from macromolecules, the modeled signal is considered GABA+macromolecules (GABA+).

**Figure 2.**
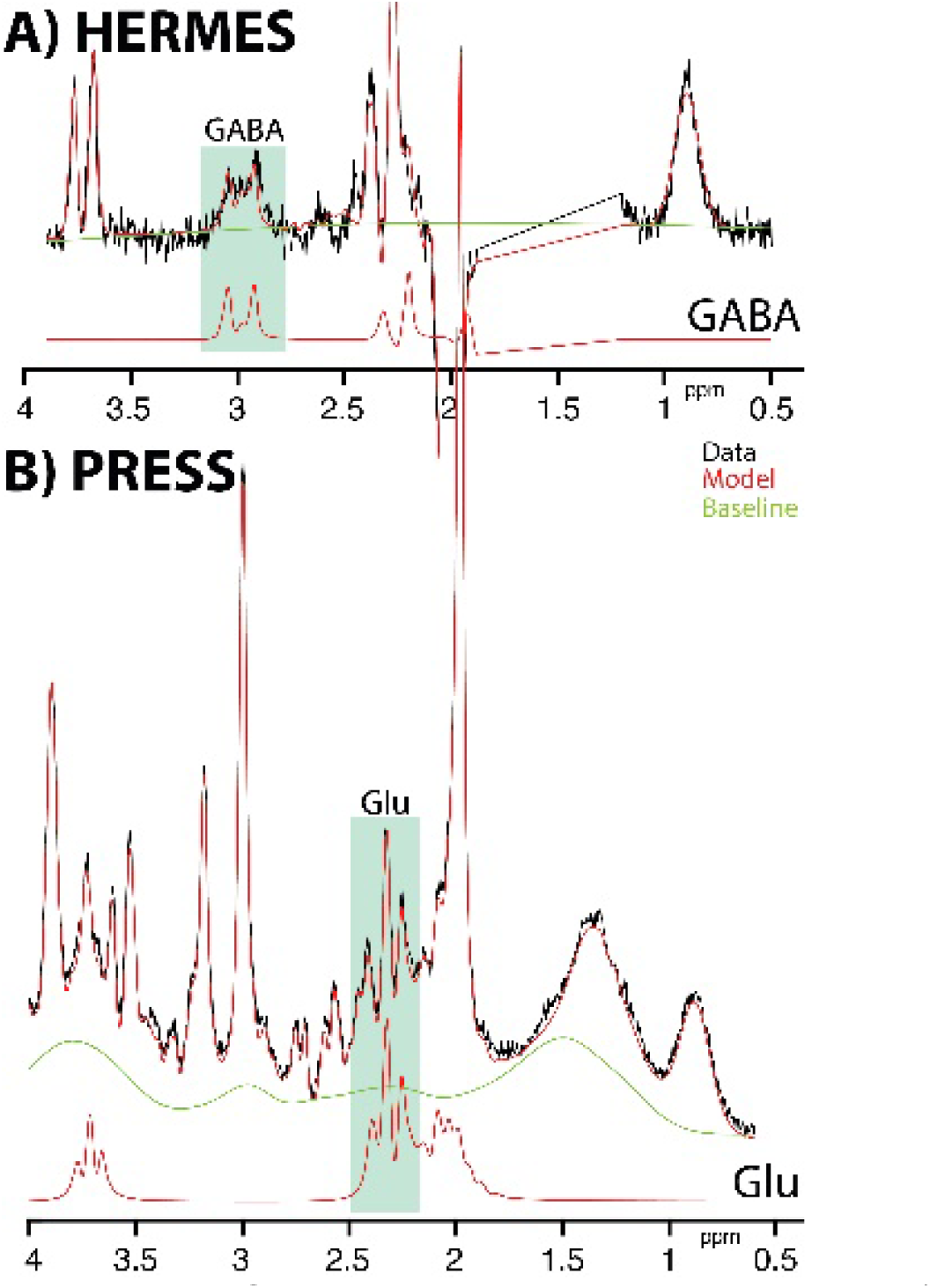
MRS spectra and respective models from A) the HERMES sequence yielding a GABA-edited spectrum with the GABA signal detected at 3 ppm, and B) the unedited PRESS spectrum with the Glu signal at ∼2.34 ppm.

### 2.8 Statistical Analyses

Statistical analyses were performed using SPSS (IBM SPSS Statistics Version 32). Independent-sample *t-*tests examined TD/ASD group differences in age, estimated nonverbal IQ (eNVIQ), and SRS-2, as well as the MEG midline parietal-occipital source exponent and the MRS midline parietal-occipital voxel Glu, GABA+, and Glu/GABA+ ratio. Although for all association analyses the primary focus was MRS chemistry and RS exponent relationships, group was included as a factor. Hierarchical regressions examined associations between MRS measures (run separately for Glu, GABA+, and Glu/GABA+ ratio) and the parietal-occipital RS exponent, with the parietal-occipital MRS measure entered first, group second, and the interaction term last and RS exponent as the dependent variable.

## 3. Results

### 3.1 Determining the Specparam Frequency Range

Analyses to determine the specparam frequency range were conducted using the sample with evaluable data. MEG RS brain imaging data and MRS brain chemistry data were obtained from 157 children (101 TD and 56 ASD), providing 176 datasets (110 TD and 66 ASD). Of these, 14 MEG datasets (11 TD and 3 ASD) could not be analyzed given amount of artifact (2 TD and 2 ASD; e.g., artifact from metal filings) or participant sleeping during the RS exam (9 TD and 1 ASD)^2^. One dataset (TD) could not be analyzed due to excessive motion during the MR and MRS scan. One dataset (ASD) was excluded given the average drift values greater than a cutoff of 5 Hz for HERMES (see below).

The effect of the specparam frequency range was assessed using the sample with evaluable MEG and MRS data. As shown in Figure 3, GABA+ and exponent associations (reported as R^2^ values) were largest in the 4 to 55 Hz window, although the pattern of findings was similar across all examined frequency ranges in both TD and ASD. Given the findings and given concerns that specparam does not do a good job estimating the exponent in the presence of periodic delta activity (Gerster et al., 2022)^3^, for each child the exponent value was computed using a 4 to 55 Hz range.

**Figure 3.**
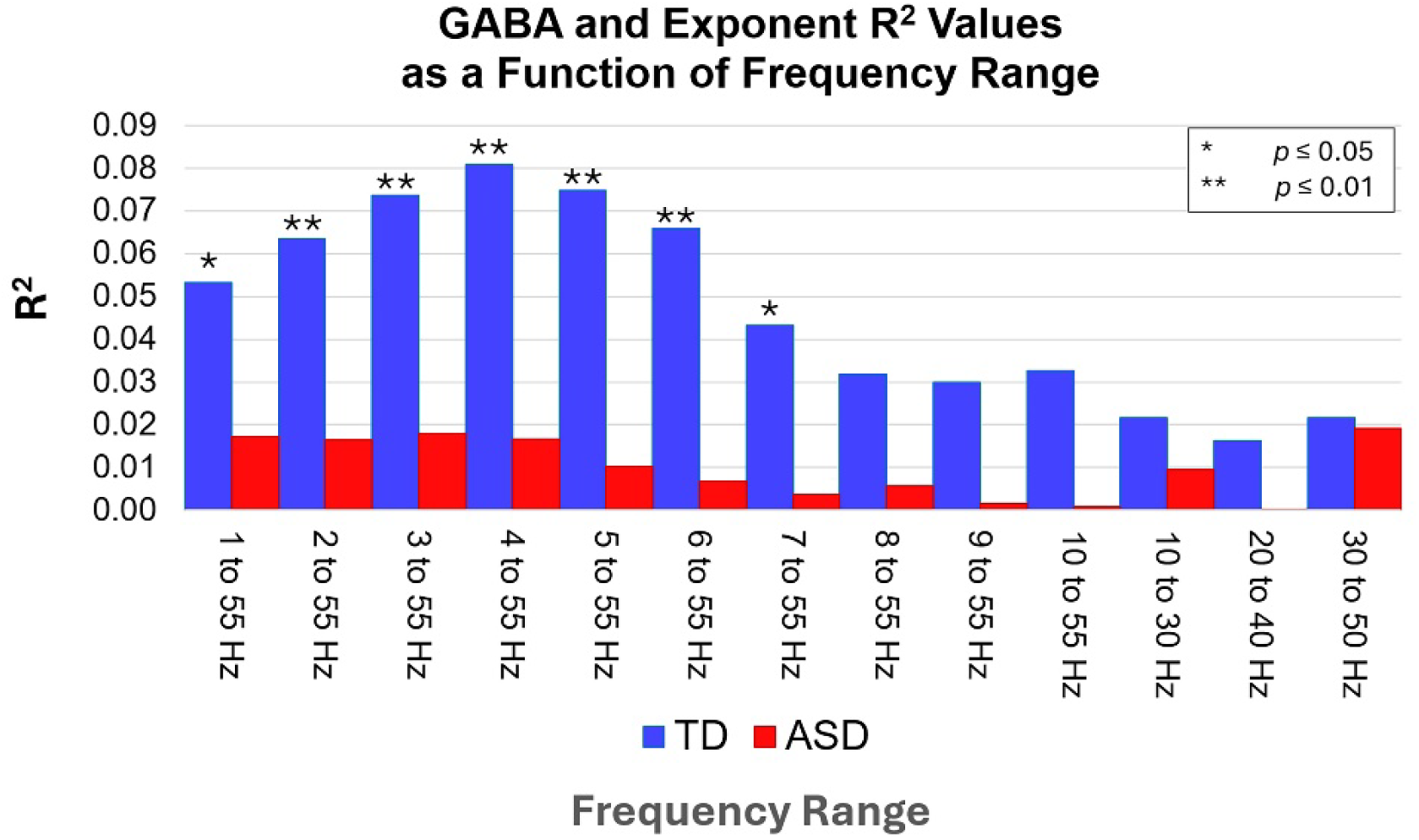
Squared correlation (R^2^) values shown for a 1 to 55 Hz range (55 Hz the upper limit given power line noise at 60 Hz) and then increasing in 1 Hz steps up to 10 Hz. For comparison with Gao et al. (Gao et al., 2017), values for 10 to 20 Hz, 20 to 40 Hz, and 30 to 50 Hz ranges are also provided.

### 3.2 Final Sample and Quality of the Data

Using the 4 to 55 Hz range, univariate and multivariate outliers were identified. Univariate outliers included 1 GABA+ value outlier (ASD), 2 Glu outliers (1 TD and 1 ASD), and 1 an exponent outlier (ASD). Eight multivariate outliers were removed (4 TD and 4 ASD). In total, 28 datasets were excluded (17 TD and 11 ASD). After these exclusions, 148 RS and MRS datasets were available from 129 children (84 TD and 45 ASD). Nineteen children (9 TD, 10 ASD) had evaluable datasets at two timepoints. Because the visits occurred more than 1.5 years apart, all available datasets were retained for analysis. Analyses excluding the second recording from these 19 children (see Section 3.4) showed patterns of findings similar to those reported below.

Analyses using the 148 evaluable datasets showed that the TD and ASD children differed in age (see Table 1; ASD children on average 1 year older). The groups did not differ on any of the IQ measures. As expected, TD and ASD children differed on SRS-2. A Fisher’s Exact Test showed that the TD and ASD groups differed in number of males and females (TD female = 47%, ASD female = 27%, *p* < 0.05). Of the combined sample, 59% were white, 16% were African American, 14% were multiracial, 9% were Asian, 0% were Pacific Islander, and 1% did not report race; 89% reported being non-Hispanic, reported being 8% Hispanic, and 3% did not report their ethnicity.

**Table 1.**

| Group | N | Mean | SD | *p-value*<sup>a</sup> |
|---|---|---|---|---|
| **Age (years)** |  |  |  |  |
| TD | 93 | 12 | 2 | 0.001 |
| ASD | 55 | 13 | 2 |  |
| **Estimated Full-Scale Intelligence Quotient**<sup>b</sup> |  |  |  |  |
| TD | 89 | 109 | 11 | 0.54 |
| ASD | 52 | 107 | 17 |  |
| **Verbal Comprehension Index**<sup>b</sup> |  |  |  |  |
| TD | 89 | 110 | 10 | 0.09 |
| ASD | 52 | 105 | 18 |  |
| **Estimated Nonverbal Intelligence Ouotient**<sup>c</sup> |  |  |  |  |
| TD | 90 | 106 | 13 | 0.44 |
| ASD | 53 | 108 | 16 |  |
| **Social Responsiveness Scale - 2nd Edition**<sup>d</sup> |  |  |  |  |
| TD | 87 | 45 | 6 | <0.001 |
| ASD | 53 | 68 | 12 |  |

<sup>a</sup>Based on a *t*-test
<sup>b</sup>Scores not avilable for 7 datasets (4 TD, 3 ASD)
<sup>c</sup>Scores not avilable for 5 datasets (3 TD, 2 ASD)
<sup>d</sup>Scores not avilable for 8 datasets (6 TD, 2 ASD)

Group differences in the quality of the MRS and MEG data were assessed. Groups did not differ in the PRESS or HERMES drift values (*ps* > 0.05). Groups did not differ in the amount of evaluable RS data (TD mean = 343 s, SD = 62; ASD mean = 346 s, SD = 48; *p* > 0.05). As shown in Figure 5 the specparam algorithm provided an excellent fit to the data from each child, with most children exhibiting R^2^ values greater than 0.97 (range 0.96 to 0.99) and with no instances of underfitting, defined as a mean absolute error (MAE) value *>* 0.10. The consistently high values indicate that the specparam power spectrum model successfully distinguished aperiodic and periodic activity. Groups did not differ in the specparam R^2^ fit (TD mean = 0.99, SD = 0.007; ASD mean = 0.99, SD = 0.006; *p* > 0.05). Although groups differed on the specparam MAE values (TD mean = 0.028, SD = 0.008; ASD mean = 0.031, SD = 0.009; *p* < 0.04), the group difference was small.

**Figure 4.**
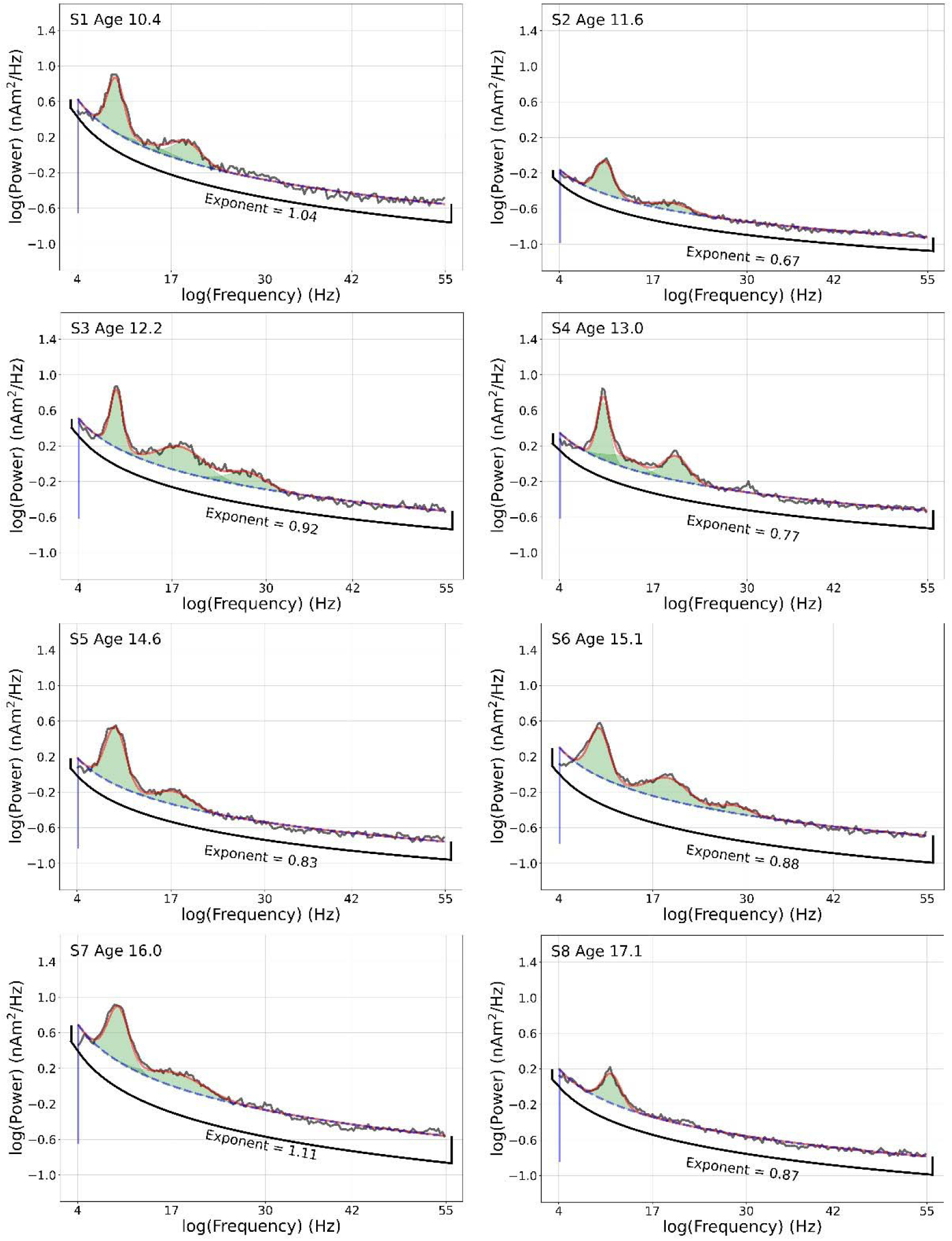
Eyes closed original power spectrum (black line) and specparam model (red line) in 8 representative children 10 to 17 years old. The x axis shows log frequency (Hz) and the y axis log power. As shown in each panel, in each child the original power spectrum (black line) is well modeled by specparam (red line). The aperiodic exponent (slope of a linear fit to the specparam model in log-log space) parameters is shown for each child.

**Figure 5.**
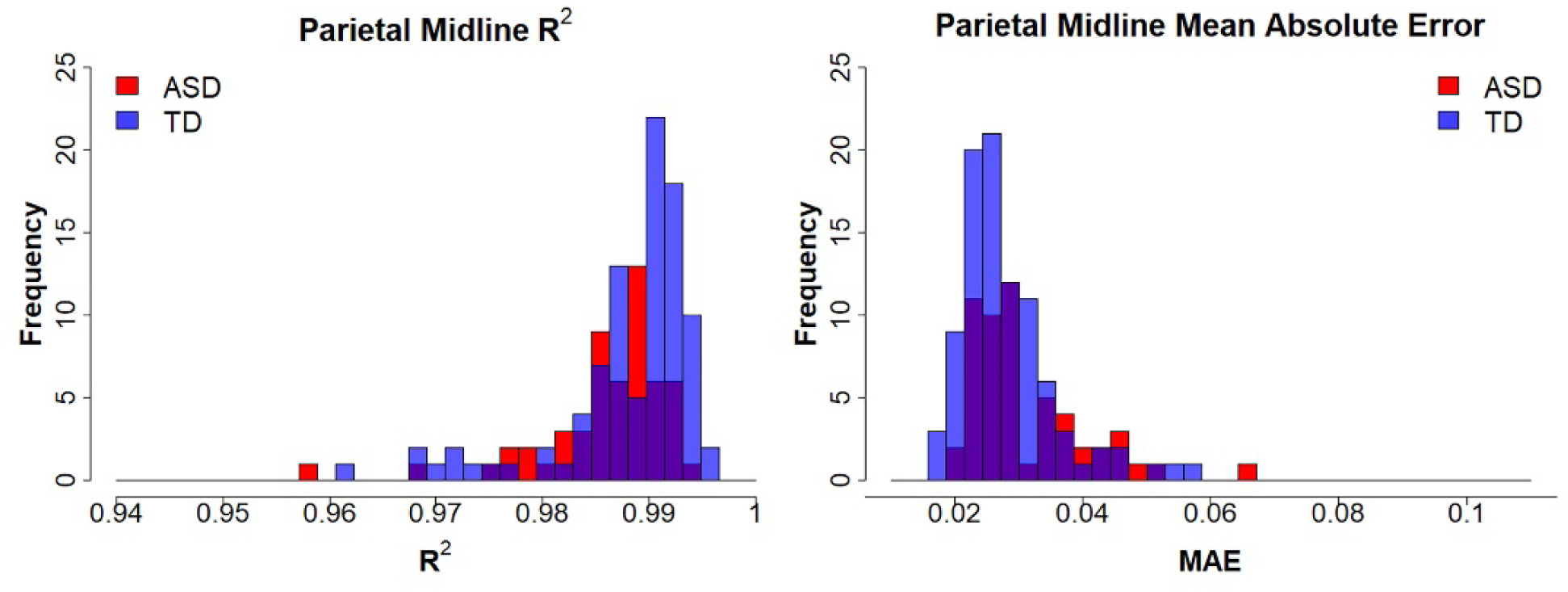
The distribution of specparam R^2^ values (left) and mean absolute error (MAE) values (right) for TD (blue) and ASD (red) groups. As detailed in Ostlund et. al. (Ostlund et al., 2022b), specparam provides two measures that describe how well the model fit the data - R^2^ and MAE. R^2^ assesses the correspondence between the original spectrum and the full model. MAE is computed examining the error between the original spectrum and the full model. Observed R^2^ values > 0.90 and MAE values < 0.08 indicate that the model fit parameters were appropriate across the examined frequency range.

### 3.3 Group and Age Effects on MEG and MRS

*t*-tests showed that groups did not differ in parietal-occipital RS exponent, Glu and GABA+ concentration, or Glu/GABA+ ratio (*p*s > 0.21). Regressions showed a marginal association between age and exponent in the full sample (*p* = 0.07; group difference *p* = 0.97). An age-related decrease in Glu concentration was observed (*p* < 0.02), with no interaction between groups (*p* = 0.93). Age was not associated with GABA+ concentration (*p* = 0.77), and groups did not differ in the age and GABA association (*p* = 0.54). Age was not associated with the Glu/GABA+ ratio (*p* = 0.39), with no interaction with group (*p* = 0.55).

### 3.4 RS and MRS Associations

Figure 6-8 scatterplots show associations between MRS (x axis) and RS exponent (y axis) for Glu, GABA+, and Glu/GABA+ ratio. No effects were observed for Glu concentration (*ps* > 0.14). Simple-effect analysis of a Group x GABA+ concentration interaction, *F*(1, 144) = 6.55, *p* < 0.01, showed a positive association between GABA+ and exponent for TD (*p* = 0.01) but not ASD (*p* = 0.40). Simple-effect analysis of a Group x Ratio interaction, *F*(1,144) = 4.28, *p* = 0.04, showed a positive association between the ratio and exponent for TD (*p* = 0.01) but not ASD (*p* = 0.85).

**Figure 6.**
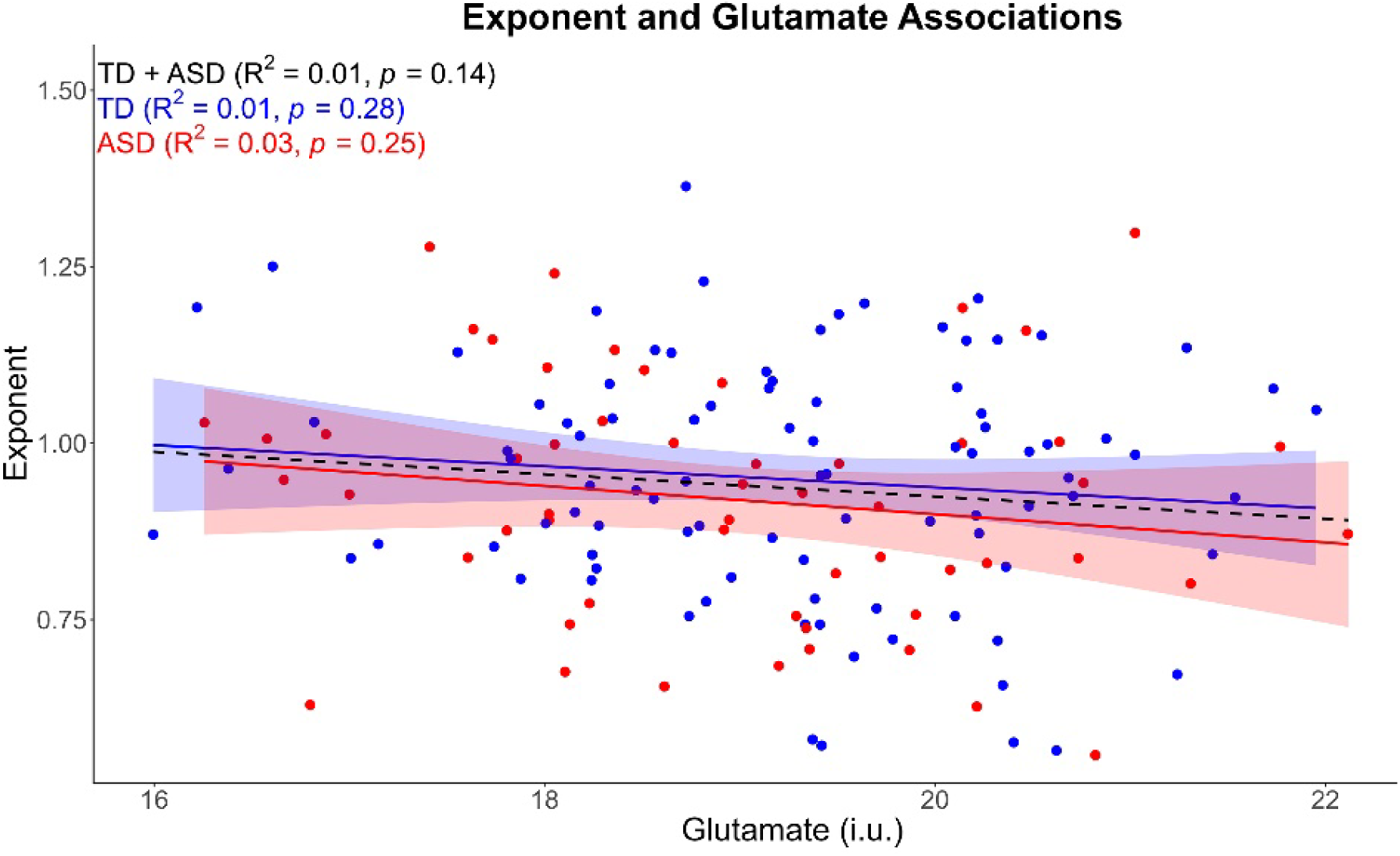
Scatterplots showing for TD (blue) and ASD (red) midline parietal associations between Glu (x axis) and RS exponent (y axis).

Regressions in the TD sample with exponent as the dependent variable and with GABA+ entered first and Glu/GABA+ second (and then vice versa) showed that neither GABA+ or Glu/GABA+ predicted significant unique variance when entered second. The pattern of all above findings remained unchanged after removing variance in the exponent associated with age (i.e., entering age in first in the hierarchical regression).

Exploratory regressions run separately for TD and ASD assessed sex differences in the above neurochemistry and exponent associations (sample size likely not large enough to detect a clinically meaningful Chemistry x Group x Sex interaction). None of the Sex x Chemistry interaction terms were significant in either group (*ps* > 0.14).

Exploratory analyses showed no differences in the neurochemistry and exponent associations using the Glu or GABA+ concentration measure that was not tissue-corrected.

Analyses excluding the second recording from the 19 children with longitudinal data showed patterns of findings analogous to those reported above. No effects were observed for Glu concentration (*ps* > 0.11). Simple-effect analysis of a Group x GABA+ concentration interaction, *F*(1, 125) = 6.27, *p* = 0.01, showed a positive association between GABA+ and exponent for TD (*p* = 0.02) but not ASD (*p* = 0.24). Simple-effect analysis of a Group x Ratio interaction, *F*(1,125) = 5.35, *p* = 0.02, showed a positive association between the ratio and exponent for TD (*p* = 0.01) but not ASD (*p* = 0.47).

## 4. Discussion

In the TD children, parietal-occipital GABA+ concentration predicted the parietal-occipital RS exponent. No such association was observed in the children with ASD, evidence of disruption of a brain chemistry/neural function association in these children. No association was observed for Glu in either group. These findings indicate that a simple method for estimating E:I ratio – one that can be applied retrospectively to existing EEG/MEG data – has the potential to remove a major hurdle in understanding a currently difficult-to-measure yet fundamental aspect of neural computation that has been proposed as important across a range of clinical conditions.

### 4.1 Association Between Brain Chemistry and Resting-State Exponent in TD Children

The observation in TD children of exponent associations with GABA+ but not Glu empirically validates Gao et al.’s (Gao et al., 2017) simulations showing associations only for GABA and the exponent. This pattern is also consistent with the simulations reported in Lombardi et al. (Lombardi et al., 2017) showing a stronger influence of inhibitory than excitatory neurons on the power spectrum exponent and with Chini et al.’s (Chini et al., 2022) mouse studies showing that strengthening of inhibition relative to excitation (medial prefrontal cortex) was associated with a larger exponent.

Comparing present brain chemistry and exponent results in the TD children to the very limited brain chemistry and scalp EEG exponent literature, McKeon et al. (McKeon et al., 2024) reported on MRS GABA concentration (prefrontal cortex) and the EEG exponent computed from anterior electrodes (right: F4, F6, F8; left: F3, F5, F7) in 164 controls 10-32 years old. They observed a positive association between Glu and the exponent and no association with GABA. They also found that more asymmetry (defined as the absolute difference in age-adjusted Glu and GABA levels) was associated with a larger exponent. McKeon et al. suggested that, “The associations with the exponent suggest that developmental decreases in Glu may be driving the decreases in Glu-GABA asymmetry, thus increasing the E/I balance which is consistent with a lower or flatter exponent.” Differences in brain chemistry and exponent associations between McKeon et al. and the present study may be due to the greater age range in McKeon et al., MRS from different brain regions, their limiting electrophysiological characterization to scalp sites (thus reflecting activity from distant brain regions), and the use of 7T MRI in McKeon et al. (7T MRI more accurately estimating Glu concentration) and 3T MR in this study.

### 4.2 TD and ASD Differences in Brain Chemistry and Resting-State Exponent Associations

The Group x GABA+ concentration interaction for the exponent measure was surprising given no group main effect in midline parietal-occipital exponent or GABA+ or Glu concentration. Gaetz et al. (Gaetz et al., 2014) also found no group difference in parietal-occipital GABA+ concentration and also that GABA+/Cr group differences were region specific (mean age of children ∼11 years old), with no group differences at a midline visual cortex ROI but with a significantly smaller GABA+/Cr measure in ASD than TD in left motor cortex (11% group difference) and left auditory cortex (22% group difference). Other MRS studies have observed that GABA group differences are regionally specific. Reporting on left and right auditory cortex GABA concentration in children with (N = 56) and without (N = 30) ASD 7 to 18 years old, Saleh et al. (Saleh et al., 2024) observed higher GABA and lower Glu concentration in ASD than TD in the right but not left auditory cortex.

Table 1 of Saleh et al. (Saleh et al., 2024) reviews many of the studies assessing Glu and GABA concentration in TD and ASD. Findings are mixed, with some studies reporting Glu and GABA group differences and others not. As Saleh et al. noted, studies report on different brain regions (e.g., anterior cingulate cortex, motor cortex, visual cortex, auditory cortex, and thalamus) and differ in participant age, method used to acquire the MRS measures, and method used to estimate Glu and GABA concentration. In general, studies to date indicate a regional specificity to the TD and ASD MRS Glu and GABA group differences.

GABA+ concentration explained a significant 8% of the variance in the RS exponent in the TD children (Figure 7), a medium effect size (Cohen, 1992), confirming the hypothesis that GABA+ and the RS exponent are linked. The present finding of no association between GABA+ and the RS exponent in children with ASD suggests group differences in the mechanisms associated with E:I balance (as assessed by the RS exponent), even though the present finding of no group difference in the RS exponent suggests that, at least at the group level, a different set of mechanisms support a ‘normal’ parietal-occipital RS exponent in older children with ASD. Exploring the neural basis of the RS exponent, Brake et al. (Brake et al., 2024) found that, in addition to GABA, biophysical parameters such as synaptic timescales and leak currents contribute to the RS exponent, with other mechanisms not explored in their study (e.g., active membrane currents and dendritic calcium spikes) potentially contributing to the RS exponent by changing postsynaptic response kinetics and amplitudes. Preston et al. (Preston et al., 2026) note that neural spike synchrony and the decay rate of a damped oscillation also influence the exponent. The mechanisms influencing E:I balance and thus the RS exponent are many and the interactions complex.

**Figure 7.**
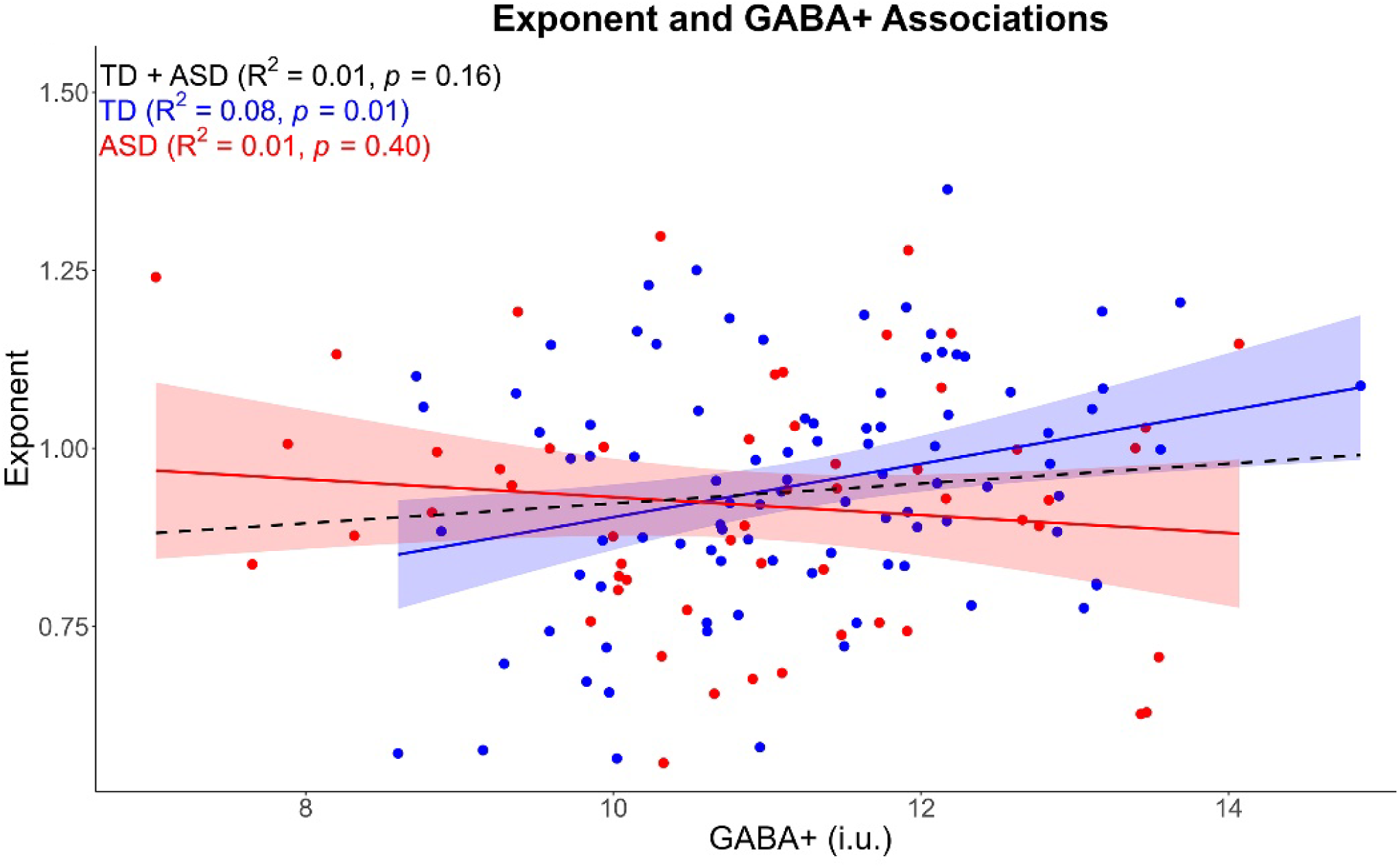
Scatterplots showing for TD (blue) and ASD (red) midline parietal associations between GABA+ (x axis) and RS exponent (y axis).

**Figure 8.**
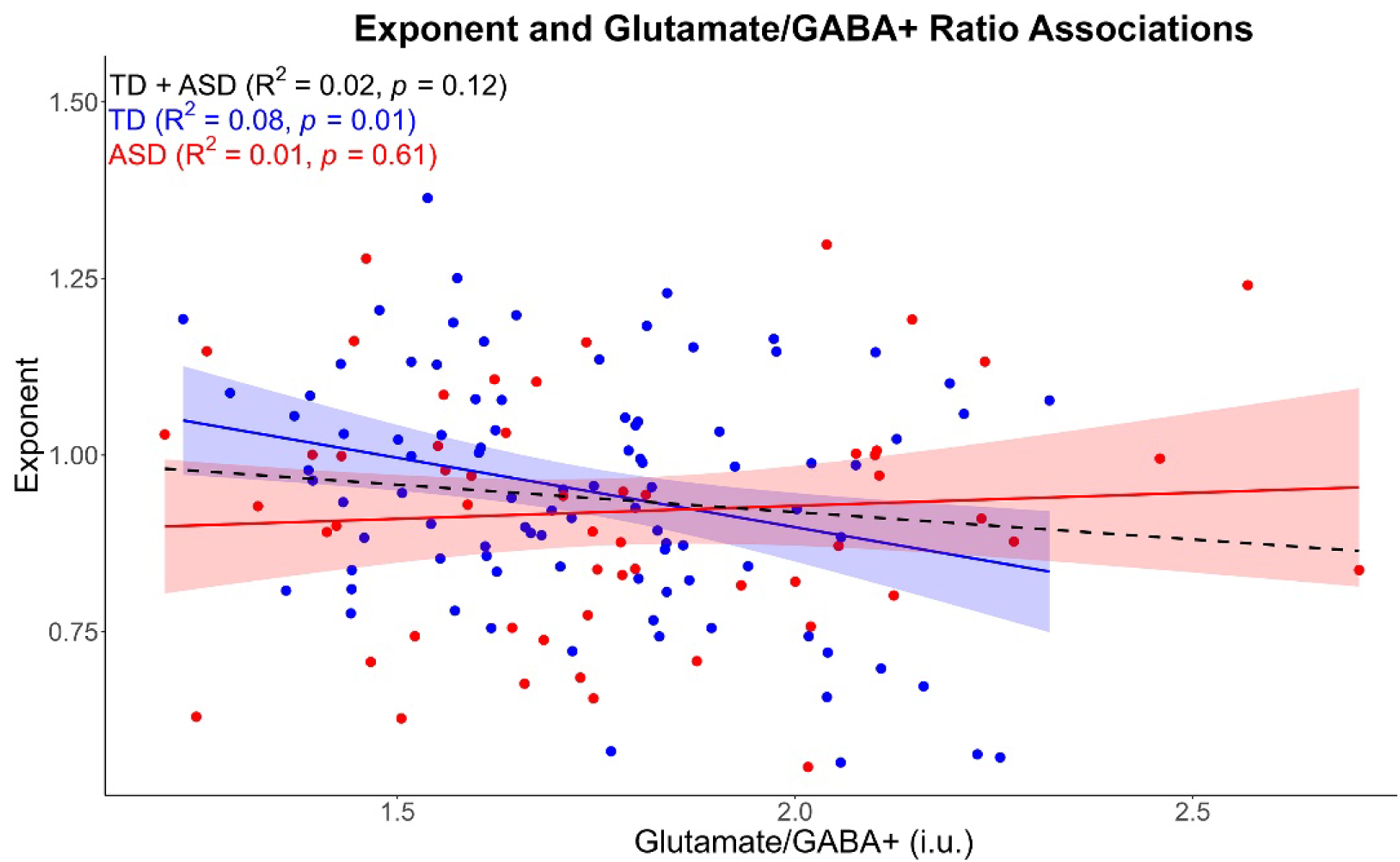
Scatterplots showing for TD (blue) and ASD (red) midline parietal associations between Glu/GABA+ (x axis) and RS exponent (y axis).

The etiology of ASD is also complex. The present study examined individuals with idiopathic ASD, which is known to be genetically complex. At the time this paper was written, the SFARI Gene database included more than 1200 genes with reported or suspected links to ASD (https://gene.sfari.org/). A definitive genetic etiology has been identified in only 25–35% of individuals with ASD (Wiśniowiecka-Kowalnik & Nowakowska, 2019). Most of the gene variations have only a small effect (Autism Spectrum Disorder Working Group of the Psychiatric Genomics Consortium et al., 2019; Fu et al., 2022; Gaugler et al., 2014; Klei et al., 2012; State & Levitt, 2011), with variations in many genes likely combining with environmental risk factors such as parental age and birth complications to determine an individual’s risk of developing ASD (Willsey et al., 2022; Yasuda et al., 2023). ASD is considered a complex, multifactorial disorder (Ruzzo et al., 2019; Treccarichi et al., 2026), with the expectation of significant between-child differences in brain development across the idiopathic ASD cohort examined in this study.

Understanding the heterogeneity in idiopathic ASD is increasingly the focus of clinical and research work, with recent studies seeking to identify subsets of children with similar features (genes, brain function measures, etc.) and thus ASD children who would benefit from a specific treatment (e.g., see Tărlungeanu & Novarino, 2018). A larger sample would be needed to identify ASD subgroups – those showing the expected GABA and exponent association versus those that do not. Looking to the future, also needed are multimodal brain imaging (MEG+MRI+MRS) pharmacological studies that modulate RS aperiodic processes (e.g., propofol – a GABAA receptor modulator that lengthens the decay time of inhibition (Kitamura et al., 2003; Orser et al., 1994)) to better understand mechanisms associated with E:I ratio and its manifestation the RS exponent.

With respect to ASD brain chemistry and exponent associations, of note is Naples et al. (Naples et al., 2026) having assessed small adult samples of ASD (N=16, mean age = 26.4 years) and TD (N=16, mean age = 27.7 years) and showing group differences in the availability of metabotropic glutamate receptor 5 (mGlu5; assessed using positron emission tomography) and in the ASD sample measuring associations between mGlu5 and the RS exponent (exponent value obtained via the average of 128 EEG sensors). Naples et al. found that across brain regions mGlu5 availability was lower by 15% in ASD than TD. In EEG from the ASD group, lower mGlu5 availability was associated with a smaller exponent at most of the examined brain regions (EEG not obtained in the TD sample). Present findings are likely to differ from those of Naples et al. given study differences in the glutamate measure (Naples et al. assessing mGlu5 receptor density versus the present study measuring extracellular and intracellular Glu concentration), their reliance on a broad average of scalp EEG (with necessarily diffuse brain sources), and study differences in participants’ ages.

Although studies comparing MRS Glu and PET Glu measures are of interest, pediatric PET studies examining Glu receptor availability are difficult to conduct given the invasive nature of PET as well as the high cost of a PET exam.

### 4.3 Associations Between Age and Brain Chemistry and the Resting-State Exponent

Although no group differences in the RS exponent were observed, present findings replicate the large and growing literature reporting an age-related decrease in the exponent (Green et al., 2025; Hill et al., 2022; McNamee et al., 2026; Rico-Picó et al., 2023; Schaworonkow & Voytek, 2021; Vandewouw et al., 2024). In the present study this age-related decrease was observed in both the TD and ASD children. Donoghue (Donoghue, 2025) reviewed the studies examining TD and ASD group differences in the power spectrum exponent and noted that the ASD versus TD exponent findings are variable – four of the studies reported a larger exponent in ASD than TD, one study a smaller exponent in ASD than TD, and five studies no difference. This variability may be due to the heterogeneity of ASD and age range differences across studies, as well as the fact that group differences are likely region-specific, and those studies were limited to scalp EEG analyses (again, with necessarily diffuse brain sources). McNamee et al. (McNamee et al., 2026) finding a significant group x age x brain region interaction for RS exponent in children with and without ASD 7 to 17years old demonstrates the need to obtain regionally-specific exponent measures. As they discussed, regional differences in the aperiodic exponent as well as in associations with age are likely due, in part, to well-known regional differences in brain maturation. Using source localization to assess RS E:I balance (assessed as functional EI (Bruining et al., 2020)) in individuals with and without ASD 6 to 32 years old, Osorio et al. (Osorio et al., 2026) found that group differences were regional and differed as a function of age. McNamee et al. (McNamee et al., 2026) and Osorio et al. (Osorio et al., 2026) findings establish the need to assess group differences in source space and for study designs to assume that group differences will change across time.

Cortex is believed to exhibit constant spontaneous aperiodic activity reflected in the exponent, with periods of synchronous firing the exception rather than the rule. The aperiodic exponent value observed for the full sample in the present study (mean = 0.94, SD = 18) is similar to that reported in Lombardi et al.’s (Lombardi et al., 2017) 1.0 to 1.14. In their simulations, this range of exponent values was associated with 20% to 30% of the synapses being inhibitory - close to that of inhibitory synapses in real neural networks (Hendry et al., 1987; Markram et al., 2004).

Whereas no association with age was observed for GABA+ in the present study, an age-related decrease in Glu concentration was observed in both groups. With respect to maturational changes in Glu and GABA, although a full review is not possible in the present paper, several main points warrant emphasis. Few pediatric studies have assessed age-related changes in glutamate. Ortuño et al. (Ortuño et al., 2025) assessed age-related changes in Glu in dorsomedial prefrontal and medial temporal lobe areas in 128 healthy individuals 7 to 34 years old. They observed an age-related decrease in Glu in dorsomedial prefrontal cortex but not medial temporal lobe. In their review of MRS studies assessing age-related changes in Glu, their Table 1 indicated mixed findings, with some but not all studies reporting adolescents showing an age-related decrease in Glu. Previous findings are difficult to compare to the present finding given that almost all previous studies assessed frontal areas (none to our knowledge reporting on parietal-occipital regions) and given differences in how Glu concentration was measured. With these caveats, present findings are generally consistent with studies reporting an age-related decrease in glutamate.

With respect to GABA, in a meta-analysis of studies including participants 8-80 years old, Porges et al. (Porges et al., 2021) showed a lifespan brain GABA trajectory in which an early period of rapid increase is followed by a period of stability during early adulthood and then a gradual decrease during late adulthood. Perica et al. (Perica et al., 2022) assessed age-related changes in the balance of GABA/creatine (Cr) in frontal cortex in 144 10- to 30-year-olds. Age-related changes in GABA/Cr were heterogenous, with a mix of stable and decreasing age effects. In the present study, lack of associations between age and GABA+ may indicate no substantial, systematic change in GABA+ concentration in parietal-occipital cortex in this age range or may be due to the relatively restricted age range of the sample (10.2 to 17.9 years).

### 4.4 Study Limitations

First, brain chemistry and neural activity associations were examined at only a single location. Studies assessing chemistry and neural activity associations in other brain regions are needed. Gaetz et al. (Gaetz et al., 2014) and Saleh et al. (Saleh et al., 2024) recommended auditory cortex as of interest in studies comparing TD and ASD. Second, the present study assessed children with ASD with relatively low support needs. Studies assessing these associations in children with higher support needs are needed to evaluate the generalizability of present findings and their clinical implications. Third, the present GABA+ measure can include overlapping macromolecule signals that can impact the GABA measure, and the MRS voxel was relatively large. Source-localization electrophysiology studies with 7T edited and unedited MRSI, providing a chemically and anatomically more precise measure of GABA and Glu (Perica et al., 2022), respectively, would be of interest.

### 4.5 Conclusion

The present associations between brain chemistry and RS exponent suggest that a relatively simple, non-invasive method for estimating E:I ratio removes a major hurdle in understanding a currently difficult-to-measure, yet fundamental aspect of neural computation that has been proposed as central to clinical conditions such as ASD, epilepsy, and schizophrenia. Present findings indicate that the contribution of local brain chemistry to neural-circuit E:I balance differs for older children with and without ASD.

## Acknowledgments

DISCLAIMER: Dr. Edgar declares intellectual property relating to the potential use of electrophysiological auditory markers for treatment planning in clinical ASD.

This study was supported in part by NIH grant R01MH107506 (JCE), NICHD grant R01HD093776 (JCE), NICHD grant R01HD119165 (YC), TALK Supplement Grant GRT-00003651 (YC), NICHD grant K08HD114880 (HLG), NIH R21DA065088 (MGS) and NIH R00DA051315 (MGS)

## Author Contributions

JCE conceptualized and designed the study. JCE, YC, MS, HLG, MM, and MP developed and implemented data acquisition and/or processing pipelines. LB and JCE developed the clinical and cognitive assessment battery. JCE, MP, MM, and DLM conducted the statistical analyses. JCE, HLG, GAM, MS, and MP made substantial contributions to the interpretation of the overall data. JCE, MD, BF, MP, AO, MM, HLG, MS, and DLM participated in data acquisition and analysis. JCE drafted the manuscript. All authors substantively contributed. All authors read and approved the final manuscript.

## Data Availability Statement

The datasets generated and/or analyzed will be made available upon completion of this study and reasonable request.

## Footnote

1. Individuals on the autism spectrum, their parents, and professionals in the field differ regarding the use of person-first (e.g., children with ASD) or identity first (e.g., autistic child) language (Kenny et al., 2016). With respect for divided opinions, both approaches to terminology are used.
2. At the start of this study, resting-state recordings were obtained at the end of the exam. Given that several children (mostly control adolescents) were falling asleep at the end of the MEG session, the order of tasks was reversed for subsequent participants, with the resting-state task administered first.
3. Gerster et al. (Gerster et al., 2022) identified features of some power spectrum density (PSD) functions that could result in specparam errors when estimating the exponent. Problematic features of the PSD include: a frequency fitting range that includes spectral plateaus (at low and high frequencies), a fitting range that includes oscillatory peaks at the lower or upper border of the fitting range (e.g., a periodic delta peak leads to a steepening of the spectrum, which leads to errors in estimating the exponent), and multiple overlapping periodic peaks.

With respect to delta activity, Gerster et al. (Gerster et al., 2022) noted that, “Overall, fitting and removing delta oscillation peaks seems unfeasible since they rarely occur as a single distinguishable peak in the double logarithmic representation. Furthermore, FOOOF [now called specparam] requires smooth input spectra to reduce the impact of noise which at the same time hinders fitting sharp peaks. Therefore, we recommend 1/f estimation for a higher lower-border of the fitting range to avoid the impact of these low-frequency oscillations. For high lower borders of the fitting range, oscillations can be better avoided, and if they are present, they likely have less impact on the estimation.” This may explain in the present study the larger GABA and exponent associations for the 4 to 55 Hz range versus fitting ranges that included delta activity. Yet in this study findings for lower frequency ranges were similar to those observed at 4 to 55 Hz. The weaker GABA and exponent associations in the 7-55 Hz to 10-55 Hz fitting ranges are likely due to the difficulty Gerster et al. noted in estimating the exponent when a periodic peak exists – in this case the alpha peak.

Illustrated in Figure 4, periodic peaks in the alpha and beta to low gramma ranges were observed. Gerster et al. noted that, “MEG and EEG cortical data of healthy participants typically have peaks in the ranges 8–13 Hz and 18–25 Hz which does not impose considerable challenges for the estimation of the aperiodic part of the spectrum.” In the present study, given a 4 to 55 Hz range, the alpha and beta periodic activity was well modeled.

